# Tree-ensemble analysis tests for presence of multifurcations in single cell data

**DOI:** 10.1101/200923

**Authors:** Will Macnair, Laura De Vargas Roditi, Stefan Ganscha, Manfred Claassen

**Affiliations:** Institute of Molecular Systems Biology, ETH Zurich, Switzerland; Current affiliation: Institute of Pathology and Molecular Pathology, University Hospital Zurich, Switzerland

## Abstract

We introduce TreeTop, an algorithm for single-cell data analysis to identify and assess statistical significance of branch points in biological processes with possibly multi-level branching hierarchies. We demonstrate branch point identification for processes with varying topologies, including T cell maturation, B cell differentiation and hematopoiesis. Our analyses are consistent with recent experimental studies suggesting a shallow hierarchy of differentiation events in hematopoiesis, rather than the classical multi-level hierarchy.

---

Many important biological processes, such as differentiation in developmental and immune biology, and clonal evolution in cancer, can be conceived of as bi- or multi-furcated cellular state trajectories. Hematopoiesis is such a process, where hematopoietic stem cells (HSCs) give rise to multiple distinct mature blood cell types via a sequence of lineage commitments. The exact sequence is still debated^1^, either assuming a hierarchical architecture of multiple fate decisions via distinct oligopotent progenitor cell states^2–4^, or a flat hierarchy of hematopoiesis, which does not include oligopotent progenitors, and where HSCs differentiate directly into committed lineages^5–7^.

High dimensional single-cell technologies, such as single-cell RNA sequencing^8^ and mass cytometry^9^, constitute increasingly widely used tools to investigate such opposing models of differentiation, and other branching processes. These technologies allow the evaluation of the state of single cells, i.e. the transcriptional or proteomic abundance profile in the case of single cell RNA sequencing or mass cytometry, respectively. Biological processes can be conceived of as trajectories through state space: temporal sequences of cellular states that can either be derived from time series or reconstructed from non-time series single-cell data^10^. We define a *branch point* as the location in state space where three or more distinct cellular state trajectories meet. Branch points dissect these trajectories into distinct state trajectory branches.

Identifying branch points is challenging because for each single cell measurement, both branch membership and ordering within each branch must be learned simultaneously. Existing approaches include pseudotime ordering, which learns a latent time variable along a mean trajectory through state space is limited to non-branching processes^11–12^. SPADE overcomes this limitation by fitting a single minimum spanning tree to non-deterministically clustered data^13^. Monocle^11,14^ fits smoothed trees to a low dimensional representation of single cell data, where branch points in the tree are assumed to correspond to branch points in the data. Both Monocle and SPADE by definition impose a tree topology, regardless of the actual topology of the data. Wishbone^15^ and diffusion pseudotime^16^ both use an embedding whose distances correspond to those along the underlying low-dimensional manifold representing the data via diffusion maps^17^. Distinct branches are then identified via anti-correlations in graph distances to a selected root point that has to be sensibly defined a *priori.* These algorithms all return a branch point regardless of the actual evidence in the data, and any decision on the presence or absence of a branch point must be made by the user.

We introduce TreeTop to address these shortcomings: critically, the inability of other algorithms to inform users whether a branch point is present, and in addition the supervised root point selection of Wishbone, and the strong topological assumptions of Monocle. TreeTop takes as input high-dimensional single-cell measurements. Via a representation of the data as an ensemble of trees, it identifies and assesses the statistical significance of branch points, which may join more than three branches. TreeTop first approximates the topology of the input dataset by an ensemble of trees (**Fig 1a**). A set of reference nodes, representing subpopulations of cells with similar states, is selected by the algorithm. These are connected by sampled trees, each of which may have different edges connecting the nodes, capturing the distribution of possible transitions between states of the underlying biological process. Secondly, each node is scored for branching by quantifying how consistently cutting each tree at that node partitions the ensemble of trees into separate branches (**Fig 1b, Online methods**).

To test whether there is evidence that a branch point is present in a dataset, we compare to a reference distribution of scores taken from data obtained from processes without branch points, estimating the probability that such a score might have occurred by chance alone. A commonly used statistical approach to estimate such probabilities is to calculate scores for permutations of the original data. However, we found that this test was unspecific and led to reportedly significant results in synthetic examples defined not to have branch points (**Supp Fig 1**). We therefore derived reference distributions from synthetic data defined to have an high level of structure, while not containing any branch points, and comparable to the considered experimental data, i.e. having the same number of observations and dimensionality as the test data (**Fig 1c, Supp Fig 2a, Online Methods**).

Alternative methods for identifying branch points always return the optimal branch point identified, regardless of whether one is supported by the data. We applied TreeTop, Wishbone and Monocle to the synthetic datasets used to define the null distribution (**Supp Fig 2 b-d**). Both Wishbone and Monocle identified three non-trivial branches in such datasets, where none is present. TreeTop’s branching score is defined with reference to these datasets, specifically to avoid such false positive branch point identifications.

**Figure 1:**
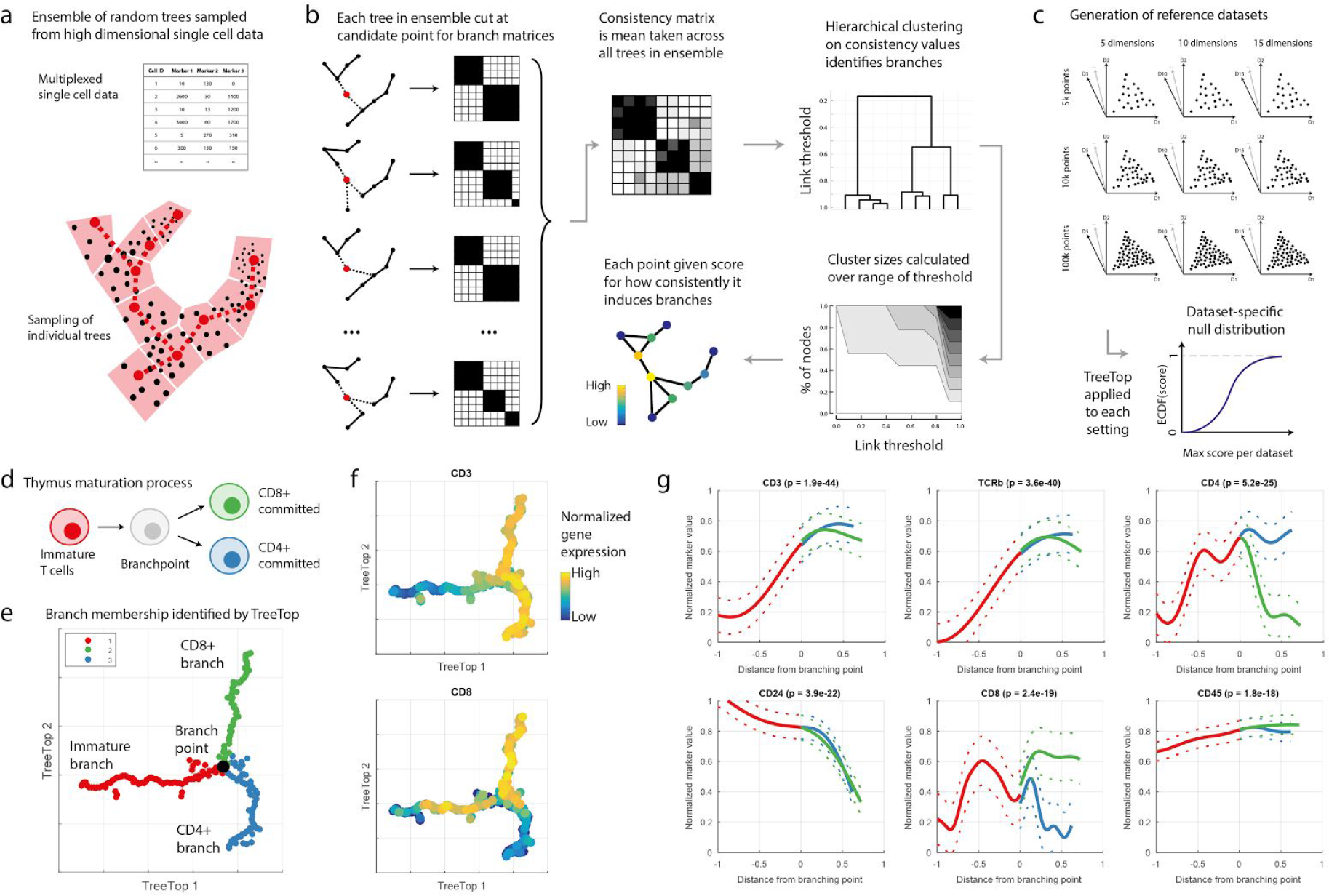
TreeTop methodology and demonstration. **a** Reference nodes are selected to be evenly distributed through data via seeding algorithm for k-means^23^; all other cells are assigned to closest reference node. Each tree is sampled by selecting a cell from each partition uniformly at random, then joining these points via a minimum spanning tree. **b** To test each reference node as a branch point (red), each tree in the ensemble is cut (removed edges are shown dashed), partitioning the remaining cells. Mean pairwise co-occurrence across all branch matrices stored in consistency matrix, i.e. (*i, j*) entry is proportion of trees in which cells / and *j* were in the same branch, when cut at the red cell. Hierarchical clustering (single linkage) performed on each consistency matrix. Sizes of the largest clusters are then calculated over all possible dendrogram cut heights, and used to score each point for branching; score is mean size of third and smaller clusters over all thresholds. **c** Branch score null distributions based on randomly generated synthetic datasets defined to contain structure but no branch points, over a range of parameters for comparison with different sizes of input datasets. **d** Cartoon of cell types in maturation of T cells in thymus. **e** Force-directed graph layout of mass cytometry thymus data (30 antibodies used^15^), pre-processed with diffusion maps (**Online Methods**)^17^. Point with highest score (black) is reported branch point, although more than one point may have a significant score. Colours indicate identified branches. **f** TreeTop layout annotated with selected protein abundances. **g** Abundance profiles for proteins on branches; x-axis is mean tree distance from identified branch point. Proteins selected for most significant differences between branches. Significance is calculated via ANOVA applied to marker abundances for each branch, Bonferroni-corrected.

Assessment of the presence of branch points has typically been done qualitatively, via visual inspection of suitable projections. However, we have found these to be misleading: the embeddings found by Monocle identify exclusively trees, regardless of the topology of the data (**Supp Fig 2d**). t-SNE^18^ projections, as used by Wishbone, are independent of the branch inference procedure, and frequently do not respect the continuity of the underlying process (**Supp Fig 2 b,c**). TreeTop visualizes the learned ensemble of trees via a force-based layout and, in contrast, provides a flexible and interpretable layout for the input dataset: comparison to the first two principal components of sample synthetic datasets shows that TreeTop’s graph-based visualization accurately captures the global structure of the data (**Supp Fig 2 e**).

**Figure 2:**
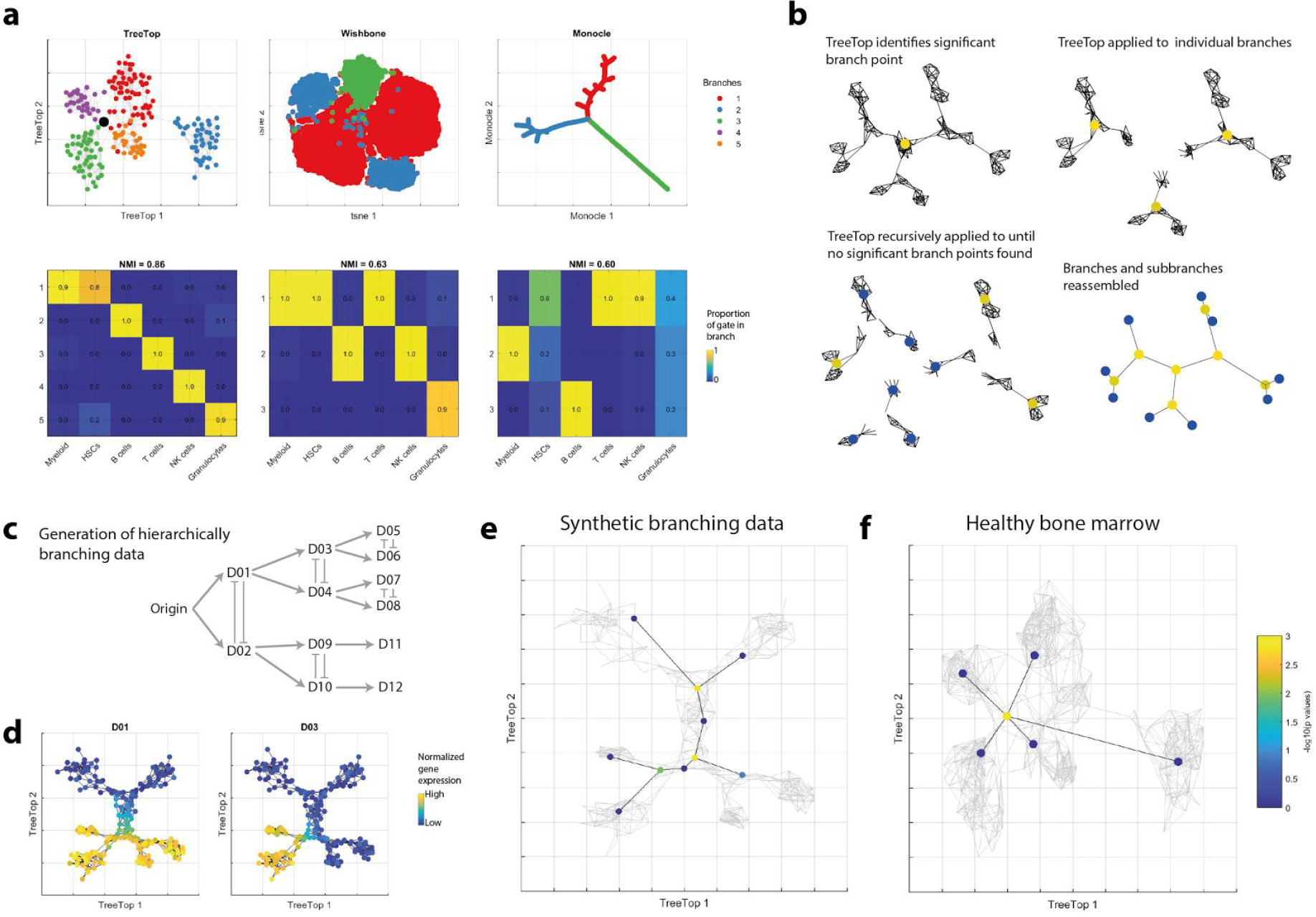
TreeTop supports shallow hierarchy model of hematopoiesis. **a** TreeTop, Wishbone and Monocle compared on healthy bone marrow mass cytometry data^19^. TreeTop and Wishbone applied to full dataset, Monocle applied to sample of 2000 cells. Top row shows layouts and branches identified by each method. Bottom row shows specificity of manual gate allocation to identified branches: heading of plot shows normalized mutual information (NMI)^24^ across all gates, branches. Ungated populations excluded. Plots showing layouts annotated by markers used are shown in **Supp Fig 9. b** Cartoon of recursive application of TreeTop to identify hierarchies of branch points. TreeTop decides whether to recurse at each step by reference to null distribution. **c** Cartoon of generation of hierarchically branching synthetic data, following classical hematopoietic architecture^2^ (**Online Methods**). **d** TreeTop layout of synthetic deep hierarchy branching data as positive control, subset of species selected to illustrate branch evolution. **e** Result of recursive application of TreeTop to synthetic branching data, showing identification of hierarchy of branch points. **f** Result of recursive application of TreeTop to healthy bone marrow data, showing only one branch point identified.

We assessed TreeTop’s capability to detect the presence or absence of branch points for biological processes with different known topologies. We applied TreeTop to mass cytometry data of T cells undergoing maturation in the thymus^15^, a process known to comprise a simple branch point with well-understood state transitions (**Fig 1d**). TreeTop assigns a p-value of less than 0.001 for the presence of a branch point. The layout shows three clearly distinct branches (**Fig 1e**), and the markers identified as showing the greatest difference between branches are consistent with biological expectations (**Fig 1f-g, Supp Fig 3**). Application of TreeTop to B cell maturation mass cytometry data^9^, a linear process, did not find any nodes with significant branch scores, consistent with the expected sequential evolution of marker abundances. TreeTop’s layout also shows a linear structure (**Supp Fig 4**). Performance of the alternative methods above was mixed, with Wishbone identifying large branches in the linear B cell maturation data (**Supp Figs 5-6**), and Monocle incorrectly ordering the B cell maturation proteins (**Supp Figs 7-8**). The flexibility of the ensemble of trees learned by TreeTop permits the accurate assessment of a wide range of topologies.

TreeTop enables us to test for the presence of branch points in data whose structure is uncertain. We assessed the two competing models for hematopoiesis, assuming either a deep or shallow hierarchy, by applying TreeTop to mass cytometry data of healthy human bone marrow^19^. TreeTop identifies a branch point (*p* = 0.002) connecting five distinct branches (**Fig 2a, Supp Fig 9);** the ensemble of trees allows branch points connecting more than three branches to be identified. Inspection of marker expression within the branches suggests that these correspond to T cells (CD3+), NK cells (CD7+), myeloid cells (CD33+), B cells (CD19+ CD20+), and granulocytes (CD24+/CD15+) (gating description in **Supp Fig 10**). These branches are connected by a branch point which does not express any markers for differentiated cell types, corresponding to the HSC compartment, consistent with its expected central position within the hematopoietic process. TreeTop enables us, in contrast to currently available approaches, to test for the presence of further subsequent branch points supporting the deep hierarchy model (**Fig 2b**). To test that TreeTop is able to find hierarchies of branch points, we applied TreeTop recursively to synthetic data specified to be representative of mass cytometry data and to include a hierarchy of branch points (**Fig 2c-e, Supp Fig 11-12, Online Methods**), and identified clear further branches. We assessed the presence of further branch points in the bone marrow data by applying TreeTop recursively to each of the 5 identified branches, and found no evidence of further branch points within the branches (**Fig 2f**). This result suggests that, within the set of markers measured, there is no evidence of any further, deeper branch points. We also applied TreeTop to single-cell RNAseq data of healthy human bone marrow^7^, and found evidence of a single branch point separating differentiated cell types into six branches (*p* = 0.001). The small branch cell counts (37-380) precluded recursive application of TreeTop (**Supp Fig 13, Online Methods**). The results from TreeTop are therefore consistent with recent findings in favour of a flat hierarchy^5–7^.

TreeTop is uniquely able to discriminate between datasets comprising up to millions of cell events for processes with and without branch points, and identify multifurcations as well as multiple levels of branch points. TreeTop provides support for the shallow hierarchy model of hematopoiesis, rather than the classical deeper hierarchy based on oligopotent progenitors. The complexity of modern single-cell data requires dedicated computational approaches such as TreeTop to identify possibly branched transitions between new cell types, and to reexamine assumed state trajectories in the light of new high-dimensional technologies, such as single cell sequencing and mass cytometry. Such approaches have led to reevaluations of long-standing previous hypotheses about the complexity and plasticity of cell types in both health and disease, including identification of new cell types such as emergency NK cells^20^ and human innate lymphoid subsets^21^, and transitions between them, such as myogenic progenitors^22^. TreeTop provides a tool based on a novel conceptual approach to test for the presence of branch points for biological processes observed via high-dimensional single cell datasets, and examine findings via an interpretable layout. We provide a MATLAB package for TreeTop.

## Acknowledgements

Will Macnair and Laura de Vargas Roditi were supported by the SystemsX.ch RTD PhosphonetPPM grant. Laura de Vargas Roditi was further supported by the ETH fellowship FEL-32 14-1 (ETH foundation).

## Author contributions

W.M. developed the method, performed the analysis and wrote the paper and the supplement. L.D.V.R. helped develop the tree ensemble sampling method, and gave feedback on the paper. S.G. simulated the synthetic branching data and wrote the relevant methodology section, and gave feedback on the paper. M.C. supervised the study, contributed to the method development and wrote the paper.

## Competing financial interests

The authors declare no competing financial interests.

## Online Methods

### TreeTop methodology

#### Overview

This section describes the geometrical intuition motivating TreeTop. branch points can be thought of intuitively as the location in some space where three or more distinct state trajectories meet. In our case, the space is the state space of cells, consisting of possible vectors of species abundances for individual cells, where species correspond to proteins in mass cytometry, and to mRNA in single cell RNA sequencing. To identify such points, we sample an ensemble of random trees representing possible transitions in the dataset, then score every point based on how consistently it partitions the remaining points into distinct branches.

Briefly, we first sample an ensemble of random trees defined over a set of reference nodes, i.e. a representative subset of cell measurements. Each tree uses the same set of reference nodes, however the connections between them may differ. In regions where data has consistent structure, many connections will be common across trees; in diffuse regions, we will observe high variability in connections. The ensemble of trees therefore captures how the subpopulations of cells may be connected to each other.

We use the learned ensemble of trees to look for branch points by considering each reference node in turn. For a given tree we cut the tree at this reference node, partitioning the tree into branches. We then compare these induced branches across all trees, looking for consistency between them: a point where branching is observed will partition the remaining points into at least 3 branches which are similar across many trees, while one with no branching will show little similarity between the branches, or less than 3 induced branches. Hierarchical clustering on the consistency matrix allows us to identify these branches, and to quantify how consistent they are.

TreeTop fits trees to the data, which capture transitions between cell subpopulation. Important implicit assumptions of our method (and the alternatives), are therefore that the data is sampled from a continuous biological process, and that the data is supported across the full range of the biological process of interest. If these criteria are not met, the cell subpopulations are separated in state space, and no evidence is available to determine the likely connections between them.

#### Data preprocessing

Mass / flow cytometry data is arcsinh-transformed (with cofactors of 5 for mass cytometry^9^ and 150, or otherwise depending on the fluorescent tag, for flow cytometry). Where the data is a mixture of non-overlapping components, the dimensionality reduction technique diffusion maps^17^ results in embeddings of the mixture components which are approximately orthogonal, with one diffusion component per mixture component^25^. This motivates preprocessing data with diffusion maps, then taking the diffusion components with the largest eigenvalues as inputs to TreeTop.

We have implemented several possible distance measures, including L1 (Manhattan distance), L2 (Euclidean) and angle distance. We have used L1, except in the case of B cell maturation data, where for consistency with the original paper we used angle distance. We have found little difference in results between L1 and L2.

#### Construction of ensemble of trees

TreeTop first selects reference nodes, which represent subpopulations with particular species expression profiles. Initially, we perform density-based downsampling of the data, to remove outliers which could cause shortcuts, and to reduce bias towards species expression profiles more densely occupied by cells (as described in Qiu et al.^26^). The user must give an appropriate value of *σ* for calculation of density (TreeTop provides functionality to assist users in this decision). TreeTop then selects a small number *k* of reference nodes, chosen to be evenly distributed through the data, thereby avoiding redundant concentrations of reference nodes in the same region. This is based on an algorithm developed for efficient initialization of k-means^23^. We use *k* = 200 throughout this study, as a balance between too few nodes, which would not allow accurate representation of all transitions in the data, and too many, requiring extensive computation for little increase in resolution. The number of nodes selected does not have a large influence on the branches identified, or their significance level (**Supp Fig 14**). The remaining cells are then labelled according to their closest reference node, partitioning the dataset into a Voronoi tessellation. Density-downsampled cells are excluded for the purpose of selecting the *k* nodes, but included for calculating the Voronoi tessellation; outliers are permanently excluded from the dataset.

TreeTop then samples a random ensemble of *n* trees connecting these reference nodes. Within each Voronoi partition, one cell is selected uniformly at random, giving the same number of points as the reference node. These are then joined by a minimum spanning tree (MST), giving a unique set of edges which connect the reference nodes and do not contain cycles. For each tree, the edges identified are recorded in an adjacency matrix, where edge weights correspond to the distances between the selected cells.

#### Ensemble of trees visualization

Visualization of the data is based on a force-directed graph layout algorithm. This class of approaches takes as input a graph, consisting of nodes connected by edges, where the edges may have weights associated with them. These can be viewed as springs, which are in a low energy state when the distance between the ends is similar to their weight, and in a higher energy state when it is different. Force-directed graph layout algorithms seek an embedding of the graph in a low-dimensional space which minimizes the resulting energy.

To apply this to TreeTop, we first take the mean over all edges in all trees in the ensemble, resulting in a graph which has an edge between two nodes where that edge occurred in at least one of the trees. Each edge has two values associated with it: the proportion of trees in which it occurred, and the mean distance between the nodes across those edges. To improve clarity of the graph, we then remove edges which occurred with low frequency, applying the maximum possible threshold which still results in a connected graph. We then apply a force-directed graph layout algorithm to this graph^27^.

#### Identification of branch points

We identify branch points by evaluating how consistently a given node partitions the other nodes into branches, across all members of the ensemble of trees. We quantify this consistency by a branching score defined as follows. If we ‘cut’ a tree by removing any of its nodes, by definition of a tree, this partitions the remaining nodes into disconnected components, or ‘branches’. Taking a node *x* and removing *x* in all *n* trees *T*_1_,…,*T*_*n*_ of the ensemble, we obtain *n* partitions of the remaining nodes into induced branches. For all pairs of nodes *i, j* ≠ *x* we then calculate the proportion *b*_*xij*_ of the trees in which *i* and *j* were assigned to the same branch. If *b*_*xij*_ is close to 1, then cutting at *x* consistently placed *i* and *j* into the same branch; if it is close to 0, then *i* and *j* were rarely placed into the same branch.

If *x* is a branch point, we expect to observe distinct groups of points, which were placed into the same branch with high probability, and placed into the branches of other groups with low probability. We quantify the extent to which each node *x* satisfies this criterion via single-linkage hierarchical clustering using the *b*_*xji*_ as a similarity measure, which we use to calculate a branching score. For a given set of similarity values, the outputs from hierarchical clustering are determined by the threshold at which the dendrogram is cut. The branching score for node x is the mean size of the third largest and larger clusters across 100 cut thresholds over the interval [0, 1]; this assesses the average size of the third largest and smaller branches of a putative branch point which induces at least 3 branches. The node with the largest branching score is the identified branch point.

#### Assessment of branch point significance

We assess statistical significance of prospective branch points by estimating p-values for branching scores with respect to a suitable null distribution. We derive null distributions from synthetic datasets where branch-like structure may be present, but which contain no branch points (**Supp Fig 2**).

We considered n-dimensional synthetic datasets comprising 100,000 points with increasing degrees of structure and measurement noise, generated to be as close as possible to a branch point, while not actually containing one. A standard Gaussian distribution has no branch point, and no branch-like structure. Linear and circular data contain branches, but no branch points. Considering a multi-dimensional Gaussian distribution as the least structured extreme, with no branch point, and a branch point connecting three branches as the most structured extreme, the continuum between these two extremes suggests a triangle as a threshold case: if any further branch-like structure were present, the data would contain a branch point.

This motivated the following test distributions, to compare performance of TreeTop and alternatives:

- Gaussian: points sampled from n-dimensional standard normal distribution;
- linear: points uniformly sampled from the interval [0, 5], with standard normal noise at σ = 0.2, then subject to a uniformly random n-dimensional rotation;
- circular: points uniformly sampled from a 2-dimensional circle of radius 0.3, with standard normal noise at σ = 0.03, centred at (1, 1,…, 1), then subject to a uniformly random n-dimensional rotation; and
- triangular: points uniformly sampled from the area inside a 2-dimensional equilateral triangle of edge length 1, with standard normal noise at σ = 0.01, with one corner at the origin, then subject to a uniformly random n-dimensional rotation.

The absolute location within the defined space does not affect TreeTop analysis, which is based on relative distances between points. Each sample of the datasets described above was generated from an n-dimensional space, where n was uniformly sampled from [6, 30]. Of these, we found that the distributions of branching scores observed for triangular data were higher than the other topologies considered by a considerable margin (**Supp Figs 1-2**). This motivated deriving null distributions from triangular data, as the most conservative distribution, which would therefore give the greatest specificity to reported TreeTop results.

We then tested the sensitivity of branching score distributions on such synthetic data were affected by four construction parameters: the standard deviation of the measurement noise added to the data (*ρ*), the number of points in the synthetic dataset (*N*), the number of dimensions in the synthetic dataset (*d*), and the number of reference nodes for the TreeTop runs (*k*). We tested the effect of each of these on the distribution of the scores (**Supp Fig 15a-e**). This analysis showed that increasing the standard deviation of noise decreases the scores identified (**Supp Fig 15a**). To have the most conservative null distribution and therefore the most specific test, we defined the null distribution by synthetic data with no measurement noise. We note that, in the context of biological data, the comparison to a distribution with zero noise is a stringent test.

We observed that the median of the branching score distribution decreases with *N,* the number of observations in the sample (**Supp Fig 15b**), and with increasing number of reference cells, *k* (**Supp Fig 15c**). Without measurement noise, number of dimensions *d* has no clear effect on null distributions (**Supp Fig 15d**), although in the presence of any noise, branching scores decrease with increasing dimensionality (**Supp Fig 15e**). To check that these trends hold for all combinations of parameters, we first generated a range of 1000 random null distributions with all possible combinations of the following parameter lists:

- Number of points: 10000, 20000, 30000, 40000, 50000, 60000, 70000, 80000, 90000, 100000
- Number of reference nodes: 50, 100, 150, 200
- Number of dimensions: 5, 10, 15, 20, 25, 30

This gave a set of 240 null distributions. We then fitted a linear model to estimate the 95th percentile of each distribution using the following input variables: 1/*N*, 1/*k* and *d*. This showed that *N* and *k* had significant reciprocal relationships with the 95th percentile values, while *d* was not found to be a significant explanatory variable (**Supp Fig 15f**). This allows us to determine, for any given run, the most conservative calculated null distribution for comparison. We therefore include in our TreeTop package this set of null distributions, and we have calculated a lookup table based on a range of numbers of observations and dimensionalities; for a given run, the package automatically selects the closest most conservative distribution for comparison.

We also considered branch score null distributions from permutations of the original data. However, tests on the basis of such null distributions yielded significant results in almost all considered real and synthetic data: permuted data contains no structure, while it is possible for datasets to contain structure without any branch points.

#### Multi-layer branch point identification

Recursive application of TreeTop can reconstruct deep hierarchies of branch points. TreeTop applied to a branching dataset identifies a branch point, and the corresponding branches induced at this point. We recursively apply TreeTop to each of the branches, resulting in branch points and, where there is evidence of a significant branch point, further branches within these (**Fig 2b**). For visualization we use the force-directed graph layout for the whole dataset (i.e. the top-level application of TreeTop), and display all significant branch points obtained via recursive TreeTop application, with leaf nodes to show branches which were not significant.

#### TreeTop Pseudocode

1. Preprocessing of data
2. Calculate density of points, based on an appropriate σ
3. Density-based downsampling, removal of outlier cell events
4. Pick reference nodes via k-means **++**^23^
5. Voronoi partition of cells according to closest reference node
6. For *k* from 1 to *n*, sample tree *T*_*k*_:

a. Uniformly at random pick one point from each Voronoi partition
b. Join these by MST
c. Record details of MST: adjacency matrix with distances, IDs of cells picked, any gating of selected cells
7. Generate force-directed graph layout embedding based on ensemble of trees
8. For every candidate branch point *x*:
  a. For every tree:
    i. Cut at this branch point, giving branches *b*(*i*) = *b*(*x*, *k*, *i*) for each point *i* ≠ *x*
    ii. Record induced branches as matrix *b*_*xkij*_ = δ_b(i)b(j)_ (i.e. 1 where in same branch, 0 where different)
  b. Take mean of *bs* across all trees *k*, giving matrix B_x_ = *B*_*xij*_ = P(*i,j* in same branch | cut at *x*)
  c. Do single-linkage hierarchical clustering using *B*_*x*_ as similarity matrix, to generate dendrogram *D*_*x*_.
  d. For each threshold *p*_cut_ = 0.01, 0.02,…, 0.99

i. Cut *D*_*x*_ at this value^1^. Cutting at a value *p*_cut_ induces a clustering of the points. When points *i,j* are in the same cluster, they have probability ≥ *p*_cut_ of being connected in the average tree, while for all *i,j* not in the same cluster, *i,j* have probability < *p*_cut_ of being connected.
ii. Calculate the sizes of all induced clusters (i.e. the number of reference nodes in each cluster).
  e. Placed in descending order, these are *N*_1_,(*p*_cut_) ≥ *N*_2_(*p*_cut_) ≥ *N*_3_(*p*_cut_) ≥…. The branching score is defined as mean third largest or smaller branch size over all cut thresholds: 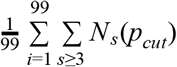.
9. Compare number of observations and number of reference nodes to null distribution lookup table, to obtain closest most conservative null distribution.
10. Compare maximum score observed with appropriate null distribution to obtain *p*-value; if significant, the reported branch point is the point with the highest branching score. The reported branches are those resulting from the threshold *p*_cut_ which gave the largest third branch.

### Generation of hierarchically branching synthetic data

As a synthetic test case with known ground truth, we simulated expression data for proteins organized in a tree of binary toggle-switches. Each switch stochastically and mutually exclusively commits to expressing one of two proteins, which subsequently activates its downstream switch and branch, respectively. Therefore one simulated trajectory mimics the multi-step differentiation process of one single cell. The structure and parameters of the underlying biochemical model were adapted from Ocone et al. (supplementary section 2.2.1)^12^.

Each protein is modelled with basal production, Hill-type functions for activation (from upstream) and inhibition (for switch) and mass-action degradation. A protein is up-regulated if activated from upstream and not inhibited within the switch:

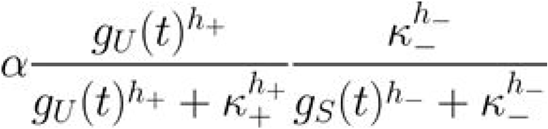

where α is the basal production rate, *g*_*U*_ is the upstream protein, *g*_*S*_ is the other protein in the switch and *κ* and *h* are the dissociation constants and Hill coefficients of the activation (+) and inhibition (−) according to **Supplementary Table 1**.

The simulation was performed with tau-leaping^28^, an approximate stochastic simulation algorithm, implemented in matLeap^29^. We simulated 1e5 trajectories with initial counts for each protein drawn from the Poisson distribution (λ = 100). The protein abundances were saved at 100 uniformly placed time points from t=0 to t=150.

#### Analysis of specific datasets

Parameter settings for each of the runs presented in this paper are described in Supplementary Table 2 for TreeTop, and Supplementary Table 3 for Wishbone.

For Monocle, all data was pre-processed via arcsinh transform, as for TreeTop. For each dataset, a sample of 2000 cells was taken uniformly at random. Monocle was run using the Gaussian distribution family, and with default values for other parameters. For the plots in this paper, the branch point which maximized the size of the smallest branch was chosen manually for each Monocle output. Monocle was first published in 2014^11^, but has since been updated to Monocle 2^30^; our analysis used Monocle 2, which for brevity we have referred to throughout the manuscript as Monocle.

#### Footnotes

1 Here we mean cutting in the sense used regarding hierarchical clustering, and not in the sense previously used for trees.

